# *‘SpikeNburst’* and *‘Nicespike’*: Advanced Tools for Enhancing and Accelerating *In Vitro* High-Density Electrophysiology Analysis

**DOI:** 10.1101/2025.02.19.638867

**Authors:** Robert Wolff, Alessia Polito, Alessio Paolo Buccino, Michela Chiappalone, Valter Tucci

## Abstract

High-density multi-electrode arrays (HD-MEAs) enable the recording of *in vitro* neuronal activity with exceptional spatial and temporal resolution. However, analysing these extensive datasets presents challenges, such as artefact removal, spike sorting, and accurate assessments of neuronal synchronization. Here, we introduce a structured protocol for conducting comprehensive HD-MEA analyses using two Python-based tools: ‘`spikeNburst`’ and ‘`nicespike`’. This protocol provides a scalable and systematic approach to processing HD-MEA recordings, ensuring efficient data handling and robust analytical outcomes.

The `spikeNburst` tool incorporates advanced methodologies for spike train filtering, burst and network burst detection, and synchronization analysis. Complementing this, we have implemented a full analysis pipeline in the `nicespike` tool, featuring GPU-accelerated spike sorting via template matching with `Kilosort`, enabling accurate identification of neuronal units across multiple electrodes. This protocol ensures more precise analyses by reducing redundancy and overestimation inherent in single-channel approaches. Moreover, both tools offer graphical user and command-line interfaces, ensuring accessibility for diverse user needs.

We validated our protocol on *in vitro* neuronal culture recordings, demonstrating its suitability to identify somatic and dendritic features of neuronal units, characterize bursting behaviour, and quantify synchronization at both unit and network levels. By addressing critical limitations of existing methods, `spikeNburst` and `nicespike` provide a robust, scalable, and user-friendly framework for HD-MEA data analysis, enhancing the study of neural network dynamics and single-cell activity in detail.

## Introduction

High-density micro-electrode arrays (HD-MEA) technologies have enhanced our capacity to study extracellular electrophysiological activity, in particular from *in vitro* and *ex vivo* systems, with unprecedented spatial and temporal resolution by capturing activity across thousands of electrodes. Furthermore, in recent years we have witnessed a growing interest in the use of HD-MEA technology as a preclinical tool in various tissue preparations, such as neuronal, cardiac, and recently of differentiated patients-derived induced pluripotent stem cells^1–3^.

Newly, progress has been made in understanding the multiple complexity scales, from single neurons to more extensive neuronal networks that characterize the nervous system. In particular, by analysing the HD-MEA recordings, it has been possible to disentangle the dynamics of a neuronal network. Furthermore, analytic methods have been developed over the years to isolate the activity of single neurons from extracellular signals. The spiking activity of individual neurons is detected with spatial and temporal information by a process called *spike sorting*^4–8^. The application of spike sorting techniques on HD-MEA data has resulted in advances in the characterization of neuronal growth, connectivity, and plasticity associated to the neuron’s physiological and pathological role in the network dynamics^2,9,10^.

However, the demand of fast, reliable, and reproducible HD-MEA results is still unmet. The complex management of the spike sorting methods is time-consuming and technically complex, resulting in limited user access and loss of data reproducibility. In addition to spike sorting, the production of high quality processing results for downstream analysis includes additional delicate steps, including artefact removal, and the accurate assessment of network synchronization^11,12^. To address these existing challenges, we provide here robust and scalable pipeline tools, namely `spikeNburst` and `nicespike,` that integrate state-of-the-art spike sorting, burst detection, and synchrony measures within an accessible and flexible framework.

### Applications

The procedures we describe are useful to enable more accurate and comprehensive investigations of neuronal activity from HD-MEAs, bridging a critical need in electrophysiological research by leveraging advanced computational methods and a user-friendly interface.

The current implementation was proven to work with electrophysiological recordings from one specific manufacturer, but it can be extended to read other file formats of raw voltage recordings, because it is open-source software and it internally depends on `spikeinterface`^7^. In particular, we adopted the HD-MEA system from 3Brain, which comprises 4,096 electrodes arranged in a square grid with an inter-electrode spacing of 60 μm. Under these conditions, neuronal activity can be concurrently detected across multiple electrodes, capturing the spatial extent of individual neuronal signals. In the protocol, we provide instructions for extending the functionality to other electrophysiological recording systems. For example, we validated that the tools can be used with HD-MEA recordings from Maxwell Biosystems devices with few changes to the source code.

Existing software solutions, including the default tools integrated within standard analytical platforms, such as the `Brainwave` software^13^ from 3Brain, primarily provide fundamental spike detection functionalities which are restricted to single-channel analyses. This limitation often results in overestimation of neuronal units and network activity due to redundant measurements of neurons being simultaneously recorded by multiple electrodes. Furthermore, while effective for systems with few electrodes, legacy tools such as `Spycode`^14^ are neither scalable nor compatible with contemporary HD-MEA datasets. This protocol addresses these gaps by presenting two Python-based tools, `spikeNburst` and `nicespike`, designed for the advanced analysis of HD-MEA recordings. Here, we describe the implementation, features, and application of these tools, demonstrating their efficacy in overcoming the limitations of existing methods while providing a scalable solution for large-scale electrophysiological datasets^15^.

### Advantages and limitations

Advantages of the tools are i) the implementation of open-source tools for the electrophysiological data analysis, ii) a ready-to-use analysis pipeline for the analysis of raw electrophysiological recordings from HD-MEA systems, iii) the integration of command-line and graphical user interfaces (GUIs) ensuring accessibility for diverse research needs from high-throughput batch analyses to individual experiment visualization. Moreover, by changing the analysis parameters in the user interface, iv) the advantage is a tool readily adapted to HD-MEA recordings obtained from a wide variety of electrophysiologically active neuronal types or tissues and v) reproducible workflow and output data easily shared.

The installation of the tools requires some basic knowledge in the Linux terminal for running the commands and working on the directory. However, the protocol provides basic computer commands that support this first step and the use of the GUIs of `spikeNburst` and `nicespike` do not require programming skills. Moreover, the type of computer used to run the analysis is a limiting factor. Sufficient working memory and CPU power is needed. A GPU supported by CUDA is necessary to run the spike sorting with `nicespike`.

Despite these limitations, the clear workflow and the data obtained by the tools allow researchers to respond to varying input datasets by choosing among the parameters in a transparent and reproducible manner and encompass comprehensive analysis at every stage of the pipeline.

## Methods implemented

We implemented multiple analyses of electrophysiological voltage recordings, summarized in Table 1.

**Table 1.** Methods implemented in nicespike and spikeNburst. ^a^These methods we re-implemented. ^b^Analyses performed with `spikeNburst` are also part of `nicespike`.

| Description | Tool | References |
| --- | --- | --- |
| Reading of electrophysiological recordings using `spikeinterface` and `python-neo` | `nicespike`<sup>a</sup> | 7,16 |
| Spike sorting with template matching using `Kilosort` | `nicespike` | 17 |
| Unit filtering using inter-spike-interval violation method | `spikeNburst`<sup>a,b</sup> | 18 |
| Burst and network burst detection | `spikeNburst`<sup>a,b</sup> | 19,20 |
| Network burst leader analysis | `spikeNburst`<sup>a,b</sup> | 21,22 |
| Synchrony: spike distance using `burst_sync` | `spikeNburst`<sup>b</sup> | 12,23 |
| Synchrony: spike-time tiling coefficient (STTC) using `burst_sync` | `spikeNburst`<sup>b</sup> | 11,12 |
| Synchrony: phase synchrony using `burst_sync` | `spikeNburst`<sup>b</sup> | 12,24–26 |
| Unit spike template characterization | `spikeNburst`<sup>a,b</sup> | 27–30 |

### Interface to electrophysiological raw voltage recordings

Spike sorting is essential for the analysis of HD-MEA voltage traces. The tool `spikeinterface`^7^ provides interfaces for different spike sorters and access to various input file formats from multiple manufacturers via the `python-neo` package^16^. For the recording system and software we used `Brainwave`^13^. There was no open-source tool for reading the raw recordings. Therefore, we implemented the reading of raw voltage traces as well as traces that have been recorded in a sparse format^31,32^. Further, the protocol uses `spikeinterface` as a python module to bandpass-filter the voltage traces (300–6,000 Hz) and to exclude noisy or broken channels.

### Spike sorting

Raw voltage traces are recorded from thousands of electrodes, also called channels, in HD-MEA systems. Because of typical small electrode distance of tens of micrometres, neuronal units can span over multiple channels. Also the sensitivity of these electrodes allows recording multiple units from single channels that may have distinct spiking features. Spike template matching is a standard procedure, well performed by `Kilosort`^17^, to circumvent these issues and to firstly detect spikes and secondly to assign them to the individual units. By default, `Kilosort` 2 is used, since it works well for static recordings and drift is minimum in *in vivo* and *ex vivo* samples. The spike sorting step is run through `spikeinterface`, which supports several other spike sorting methods and thus would make it easy to swap in a different algorithm if needed.

### Unit filtering

Additionally to filtering units by setting a minimum level of firing rate (MFR), quality assessment of units was implemented using the inter-spike-interval (ISI) violation method^18^. This method allows decreasing the likelihood of false negatives for inadequately sorted units.

### Unit characterization

Different parts of the same neuron have different spike features^27^. We implemented the characterization of units depending on the unit spike template, which is the mean of all voltage traces at a spike within a time window of 1 ms before and 2 ms after the spike. From the shape of the spike template we estimated the mean amplitude and duration of spikes for a unit. By amplitude analysis, we classified soma-like or dendrite-like units’ signals depending on the spike template shape of the central channel that has the highest voltage difference. Spike templates where the absolute minimum voltage is greater than the maximum voltage are classified as soma-like with an associated negative template height, while dendrite-like units have a positive template height greater than the negative one (cf. ‘Anticipated results‘ section). The mean spike duration, also known as peak latency, was applied to identified regular-spiking and intrinsically bursting neurons. The unit spike duration parameter, coupled with optical imaging analysis, was previously applied to identify in a culture putative glutamatergic pyramidal neurons, which show broader spikes, and putative GABAergic interneurons, which show fast spikes^28–30^.

### Burst, network burst and burst leader detection

Different firing properties and excitability of neurons significantly contribute to neural connection and information processing in the network^33^. Bursting and network bursting are complex and organized cellular behaviours that, in *in vitro* neuronal cultures, can occur spontaneously and characterize the network with brief periods of high firing rates^34^. Within a neural network, high-specialized bursting neurons act as leaders, firing at the beginning of each burst and triggering and driving the network population activity^21,22^.

We re-implemented previously described methods that relied on Matlab code (`Spycode`^14^) in python to enhance the ability to characterize burst and network burst activity. Units are bursting when they have many spikes in short time periods. A simple burst detection^19^ using only two parameters, the maximum ISI for spikes and minimum number of spikes (Table 2, Bursts), and a more advanced method with an approach for self-adapting parameters^20^ were implemented. Further, we implemented a hybrid method that uses the calculation of the ISI thresholds by the advanced method, but the burst detection with these thresholds as maximum ISI according to the simple method.

**Table 2.** Adjustable parameters. ^a^Only in `spikeNburst`, ^b^only in `nicespike`.

| Parameter | Default | Description | References |
| --- | --- | --- | --- |
| **General** |  |  |  |
| Number of parallel processes | 1 | The number of parallel processes, if 1, then no parallelization is used. | – |
| Export format | ods | The file format used for the tabular export if 'ods' or 'xlsx'. If 'pkl' then the analysis object is dumped to a file using the `pickle` module. | – |
| Analysis folder | output | The program's internal output folder used to store the analysis outputs. | – |
| **Spikes / Active units** |  |  |  |
| Use spike-sorted units<sup>a</sup> | true | Whether to use spike-sorted units instead of chip electrodes / channels. | – |
| Common active channels or units<sup>a</sup> | false | Whether common active channels/units shall be defined on the selected items. If 'all' ('any') then channels/units are active which have firing rate ≥ minimum in all (any of) selected items. | – |
| Minimum firing rate | 0.05 Hz | The minimum firing rate to define the active units. | – |
| Use valid channels only<sup>a</sup> | true | Whether to use only the valid channels saved in the `bxr` file. | 31 |
| Channel group<sup>a</sup> |  | The name of a predefined group of channels inside the bxr file which shall be used instead of all channels. If the specified group does not exist or the string is empty then all channels are used. | 31 |
| Maximum ISI violation ratio | 0 | The maximum allowed value for the inter spike interval (ISI) violation ratio. If '0' all units are selected as active. | 18 |
| ISI violation threshold | 1.5 ms | The threshold for the calculation of the ISI violation ratio. | 18 |
| **Synchrony** |  |  |  |
| STTC: time window | 0.5 s | The time window used in the calculation of the spike time tiling coefficient (STTC) for each spike $\pm \Delta t$. | 11,12 |
| Spike distance: number of bins | 120 | The number of bins used in the calculation of the spike distance. | 12,23 |
| **Bursts** |  |  |  |
| Burst detection method | pasquale | The method to use for the burst detection. The 'chiappalone' algorithm is described by Chiappalone et al. (2005) and the 'pasquale' algorithm by Pasquale et al. (2010). 'chiappalone_isi_threshold' is a modified version of 'chiappalone' using the ISI thresholds calculated as in 'pasquale' algorithm. The '3brain' method<sup>a</sup> reads previously calculated bursts by the Brainwave software. | 13,19,20 |
| Minimum number of spikes | 5 | The minimum number of spikes to form a burst. | 19,20 |
| Maximum ISI | 0.1 s | The maximum inter-spike interval (ISI) for spikes in one burst for 'chiappalone' and 'chiappalone_isi_threshold' algorithms. | 19 |
| Maximum ISI threshold | 1.0 s | The maximum allowed ISI threshold for 'pasquale' algorithm. If for some channels the calculated ISI thresholds are greater, then fallback for these channels to the 'chiappalone' method. | 20 |
| ISI threshold parameter | 0.1 s | The parameter for the algorithm choice for 'pasquale' algorithm. | 20 |
| Maximum ISI for intra burst peak | 0.1 s | The maximum allowed ISI for the intra burst peak for 'pasquale' and 'chiappalone_isi_threshold' algorithms. | 20 |
| Bin size of logarithmic ISI histogram | 0.1 | The bin size for the logarithmic ISI histogram (with base 10) for 'pasquale' and 'chiappalone_isi_threshold' algorithms. | 20 |
| Window length for ISI histogram filtering | 3 | The window length for the filtering of the ISI histogram for 'pasquale' and 'chiappalone_isi_threshold' algorithms. | 34 |
| Order of polynomial for ISI histogram filtering | 1 | The order of the polynomial for the filtering of the ISI histogram for 'pasquale' and 'chiappalone_isi_threshold' algorithms. | 20 |
| Minimal distance for peak finding | 2 | The minimal distance in samples between neighbouring peaks for finding the peaks in the filtered ISI histogram for 'pasquale' and 'chiappalone_isi_threshold' algorithms. | 20 |
| Minimum void parameter | 0.7 | The minimum void parameter for 'pasquale' and 'chiappalone_isi_threshold' algorithms. | 20,37 |
| **Network bursts** |  |  |  |
| Network burst detection method | pasquale | The method to use for the burst detection. The 'chiappalone' algorithm is described by Chiappalone et al. (2005) and the 'pasquale' algorithm by Pasquale et al. (2010). | 19,20 |
| Minimum number of units | 5 | The minimum number of units involved in a network burst | 19,20 |
| Maximum IBEI | 0.2 s | The maximum inter-burst-event interval (IBEI) used to separate intra network burst IBEIs from IBEIs between network bursts. | 19,20 |
| Minimum separation | – 0.01 s | The minimum separation of the network bursts for merging after network burst detection. If less than zero, network bursts are not merged. | – |
| Maximum IBEI for intra network burst peak | 0.2 s | The maximum allowed IBEI for the intra network burst peak for 'pasquale' algorithm. | 20 |
| Bin size of logarithmic IBEI histogram | 0.1 | The bin size for the logarithmic IBEI histogram (with base 10) for 'pasquale' algorithm. | 20 |
| Window length for IBEI histogram filtering | 3 | The window length for the filtering of the IBEI histogram for 'pasquale' algorithm. | 20 |
| Order of polynomial for IBEI histogram filtering | 1 | The order of the polynomial for the filtering of the IBEI histogram for 'pasquale' algorithm. | 20 |
| Minimal distance for peak finding | 2 | The minimal distance in samples between neighbouring peaks for finding the peaks in the filtered IBEI histogram for 'pasquale' algorithm. | 20 |
| Minimum void parameter | 0.5 | The minimum void parameter for 'pasquale' algorithm. | 20,37 |
| **Plotting** |  |  |  |
| Plot format | pdf | The file format used for the plot saving, e.g. 'png', 'pdf', 'svg', ... | – |
| Plot settings file |  | The file with additional settings for plotting. | – |
| Color map | viridis | The name of a color map known to matplotlib. Possible values are e.g. 'rainbow', 'Reds', 'cividis' and 'viridis'. | – |
| AFR bin size | 5 s | The bin size for the average firing rate in seconds. | – |
| ABR bin size | 10 s | The bin size for the average bursting rate in seconds. | – |
| ANBR bin size | 20 s | The bin size for the average network bursting rate in seconds. | – |
| Plot probe map and unit templates<sup>b</sup> | true | Whether to plot the probe map and unit templates with spikeinterface. | 7 |
| Plot wave forms<sup>b</sup> | true | Whether to plot the single-unit wave forms with spikeinterface. | 7 |
| Plot single channels<sup>b</sup> | true | Whether to plot the single channel measures with spikeinterface. | 7 |

Networks burst are subsequently defined as parallel bursting of multiple units and their detection is defined analogously. To adapt to the high number of electrodes in HD-MEA devices, we replaced the parameter of the minimum percentage of recording electrodes involved in a network burst by an absolute minimum number of spiking units. Additionally, we implemented an algorithm to merge network bursts that are overlapping or have small intervals between end of one and start of another network burst.

### Synchrony

Alternative to studying the bursting behaviour of single units, various synchrony measures have been proposed that measure relations of spike trains of two units. We used the implementation of the `burst_sync` package^12^ (with minor fixes for latest `python` versions) to calculate the synchrony of two units. First, the spike distance^23^ is a measure that is close to zero for synchronous firing. In the used implementation it includes only one parameter, which is the number of bins for the dissimilarity quantification between spike train (Table 2, Synchrony). Second, the spike-time tiling coefficient (STTC)^11^ was introduced with certain necessary and desirable properties in mind, such as symmetry, robustness for firing rate variations, recording duration and small parameter variations, etc. There is one parameter, the time window Δ*t* that can be adjusted within the proposed tools (Table 2, Synchrony). Third, the phase synchrony and the related global synchrony^24–26^ is a measure of synchrony that does not imply the use of any parameter. It has been demonstrated that different synchrony measures characterize different aspects of synchrony of spike trains^12^.

## Experimental design

### Overview of the procedures

We introduce two independent procedures (Fig. 1), describing the usage of the GUIs of i) `nicespike` (Procedure 1) and ii) `spikeNburst` (Procedure 2). The choice of procedure depends on the input data to be processed. While spike trains are directly examined by the `spikeNburst` tool, raw voltage traces are processed by the `nicespike` tool that ultimately plots and exports the analysis results for each recording using internally the `spikeNburst` tool as python module. Both tools are implemented with a graphical user interface (GUI) and can be accessed programmatically as python modules.

**Fig. 1.**
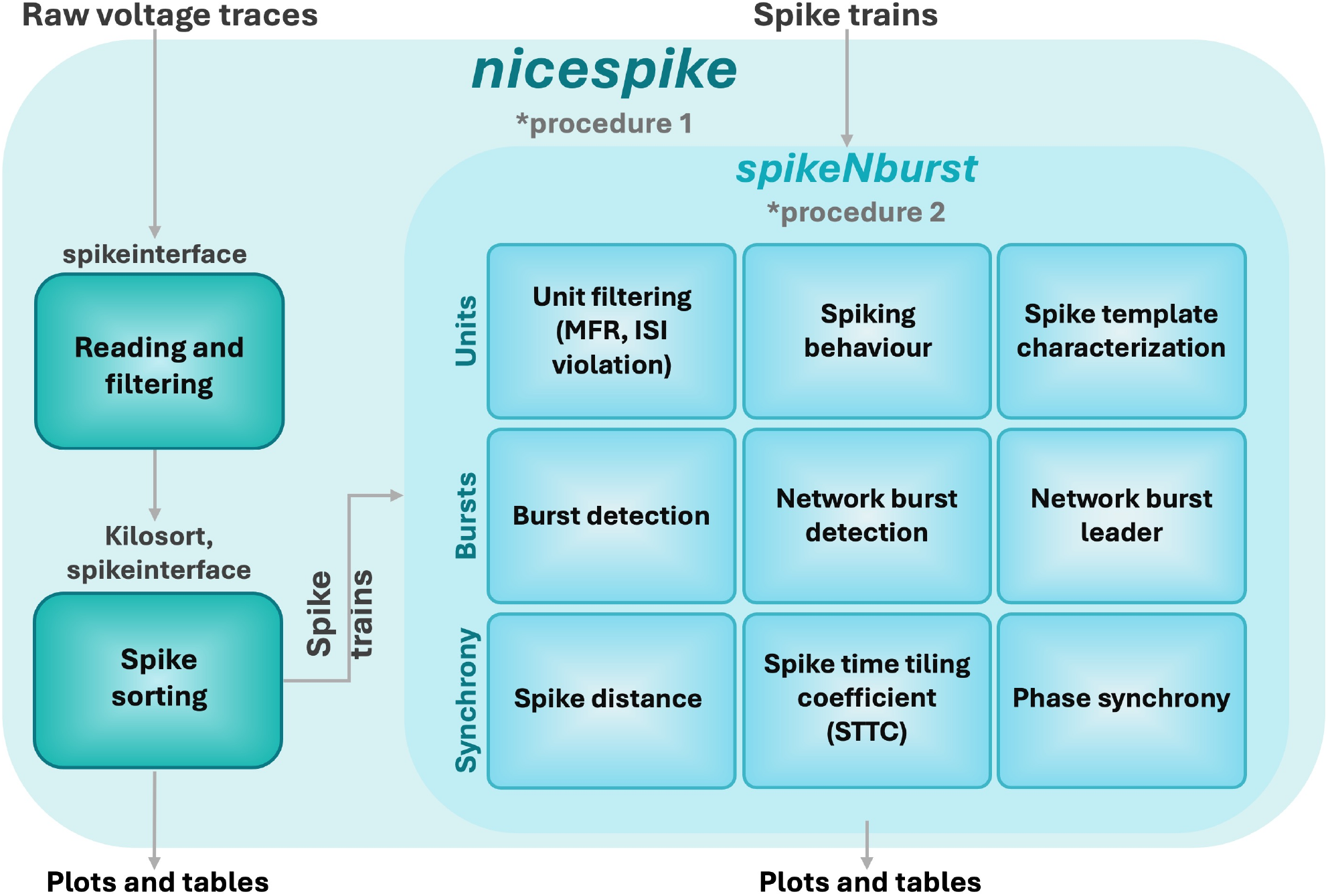
Overview of the experimental design. Raw voltage traces are analysed by `nicespike` (Procedure 1) and spike trains by `spikeNburst` (Procedure 2).

We implemented an analysis pipeline including spike sorting with template matching and subsequent spike train analysis within the `nicespike` tool. We used the `spikeinterface` package^7^ for accessing the `brw` files. For this, we co-authored the `brw` file reading in the `python-neo` tool^16^. Firstly, the raw voltage traces of electrophysiology recordings are filtered. We implemented an automatic channel removal for broken or very noisy channels that removes signals if they have constant consecutive values for at least 10 samples or if their standard deviation (SD) is > 1.7 times the median of SD of all channels. Broken channels, determined from previous recordings, are removed, and the raw traces are band-pass-filtered from 300 to 6,000 Hz. Then, spike sorting is done using `Kilosort`^17^, running on GPU. The spike-sorted unit spike traces are then exported to custom `npz` files that are further analysed by the `spikeNburst` tool. The `nicespike` tool can be used with a GUI (Procedure 1) or as a `python` module (BOX 3).

The `spikeNburst` tool was developed to provide accessible and advanced analysis of spike trains. Our demonstration supports input from both `bxr` files generated by the `Brainwave` software (3Brain, BOX 1) and custom `npz` files (`NumPy` data archive format, used internally by `nicespike`), but can be adapted to read other formats of spike train data (BOX 2). The tool incorporates multiple analytical methods. First, neuronal units can be filtered using the inter-spike-interval (ISI) violation method^18^. Second, methods for burst and network burst detection are implemented, ranging from basic approaches^19^ to advanced algorithms^20^, enabling detailed investigation of bursting behaviour. Third, a variety of unit synchrony measures are available, including spike distance^23^, the spike-time tiling coefficient (STTC)^11^ and phase synchrony^24–26^, implemented using the `burst_sync` package^12^. Parameters calculated by `spikeNburst` can be easily exported as tabular data for further analysis and visualization, and basic plotting functionality is included. The `spikeNburst` tool can be utilized both as a GUI tool (Procedure 2) and as a `python` module (BOX 4).

### Example dataset and input file formats

In the procedure, we provide instructions to enable user-friendly analysis of single-neuron activity and network dynamics from large-scale electrophysiological datasets. We chose as illustrative example a dataset obtained by recording spontaneous electrophysiological activity of WT mouse differentiated cortical neurons with an HD-MEA system (BOX 1).

#### BOX 1

**Sample preparation and recording of the example dataset**

1. We cultured mouse embryonic stem cell (mESC) line E14Tg2a (E14) on laminin-coated dishes (Sigma) in DMEM (Dulbecco’s Modified Eagle Medium, Invitrogen) supplemented with 15% ESC-certified FBS (Invitrogen), 0.1 mM nonessential amino acids (Invitrogen), 1 mM sodium pyruvate (Invitrogen), 0.1 mM b-mercaptoethanol (Sigma), 50 U/ml penicillin/streptomycin (Invitrogen), and 103 U/ml LIF (ESGRO Millipore) as previously described^35,36^. All mESC lines were regularly controlled to exclude the presence of mycoplasma using the MycoSPY kit (Duotech/Biontex).
2. *In vitro* 2D *corticogenesis* was performed by adapting a protocol previously described^35,36^. The day before the differentiation, the mESC line was plated at low density on laminin-coated dishes. The day after, the cells start to be cultured in chemically defined default medium DDM supplemented with B27-without vitamin A (Invitrogen). From day 0 to 8 of culture, the DDM medium was supplemented with the dorsomorphin analog DMH1-HCl at 1 μM (Tocris) instead of cyclopamine. At 12 days of 2D *in vitro* corticogenesis, E14 neuroprogenitor cells were plated on 0.1 mg/ml PEI (Sigma)-33 μg/ml laminin (Sigma) coated Accura (3Brain) chips according to the manufacturer’s instructions (3Brain), differentiated until 21 days *in vitro* (DIV 21), and maintained in neurobasal medium for 21 days. The cells were cultured and differentiated at 37 °C under an atmosphere of 5% CO2.
3. The recording chips (Accura, 3Brain) comprehend 4,096 micro-electrodes with an active area of 3.8 mm × 3.8 mm. At two weeks (DIV 21+14) and at three weeks (DIV 21+21) after the end of the differentiation, we firstly acclimatized the culture in the system (BioCAM DupleX, 3Brain) for 2 minutes then we recorded for 5 minutes the spontaneous activity (`Brainwave`, version 4, 3Brain). The raw voltage traces (`brw` files) were then analysed by the `Brainwave` software to detect spikes and bursts (`bxr` files) for the comparison analysis (cf. Anticipated results).

Raw voltage traces for 4,096 electrodes in a square arrangement were recorded (`brw` files, HDF5-based format containing raw data) and analysed with the manufacturer’s software for comparison (`bxr` files, another HDF5-based format containing analysis data). The `bxr` were subsequently analysed using the `spikeNburst` tool for advanced spike train analyses. Raw voltage traces from the `brw` files were processed using the `nicespike` tool, which integrates filtering, spike sorting, and subsequent analysis with the `spikeNburst` tool.

The tools can be adapted for recordings from different HD-MEA recording systems. As an example we present the necessary source code changes in `nicespike` to be able to analyse electrophysiological recordings from Maxwell Biosystems HD-MEA devices (BOX 2).

#### BOX 2

**Adaptation for inputs from different recording systems**

Both `nicespike` and `spikeNburst` can be adapted for using different input.

- To the `nicespike` tool new formats for raw voltage recordings can be introduced by changing the way how input files are read in the function `SpikeSorting.read_and_filter` in `src/nicespike/spike_sorting.py`, lines 110–186. Many file formats are already supported by the `spikeinterface` that is internally used here^7^. Also, the probe layout might need to be adapted in lines 81–85 and the plotting functions in `src/nicespike/plotting_helpers.py` may need some adaptations, too. Further, the filtering for files with the `brw` extension in `src/nicespike/ main .py`, lines 50–51, needs to include the extensions of the new file format. The `spikeNburst` module might also need some adaptations if the probe layout does not correspond to 64 times 64 electrodes, namely in the 2D plotting function `plot_2d` in `src/spikenburst/plot.py`, lines 206–258.
- To add new spike train file formats to `spikeNburst`, some adaptations must be done to the `SpikeAnalysis. init` function by adding the reading of new formats in `src/spikenburst/spike_analysis.py`, line 212. Further, for the `spikeNburst` GUI, new file extensions must be added to the filter in `src/spikenburst/gui/window.py`, line 628.
- Some parameters are hard-coded, in particular for the filtering and reading of the raw voltage recordings, such as the bandpass filtering and the channel filtering. They may be adapted by changing `src/nicespike/spike_sorting.py`, lines 36–39.
- Additionally, because `spikeinterface` supports many spike sorting algorithms, the sorting may be modified in `src/nicespike/spike_sorting.py`, lines 188–227.

An example adaptation of `nicespike` for the reading of HD-MEA recordings from Maxwell Biosystems can be found in the branch `read_maxwell` of the `nicespike` source repository.

### Adjustable parameters

The parameters of the various analyses that can be adjusted using the GUIs are described in Table 2.

## Materials

### Data

- Electrophysiological recordings as input to `nicespike`: `brw` files in versions 3.x from `Brainwave` 3–4^31^ and 4.x from `Brainwave` 5^32^ with full raw or sparse output as input to `nicespike`.
- Pre-analysed spike and burst trains as input to `spikeNburst`: `bxr` files in version 2.x from `Brainwave` 3–4^31^, with detected spikes and optionally bursts as input to `spikeNburst`; or `npz` files that are internally used by the `nicespike` tool and can be exported from it.
- The possible implementation of alternative file formats and inputs from other recording systems is described in BOX 2. 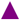 **CRITICAL** Some knowledge of python development is needed to adapt the tools to read additional file formats.
- Example datasets, which were used in the Anticipated results, are available at G-Node: https://gin.g-node.org/spiky/Wolff_et_al_2025_spiky_data (doi:10.12751/g-node.zzzzzz).

### Equipment

- Hardware: computer (test set-up: Precision 5820 Tower Workstation, Dell; Xeon W-2235 CPU, Intel; 256 GB RAM) with CUDA-accessible GPU (test set-up: Quadro RTX 5000, Nvidia). 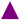 **CRITICAL** A GPU with CUDA support and sufficient memory (> 100 GB) and disk space (∼ 50 GB per 5-min recording) is needed to do the spike sorting with `nicespike`. The spike analysis done with `spikeNburst` does not use a GPU and can run on less powerful computer.
- Operating system: Linux (test set-up: Ubuntu 20.04.6; kernel 5.15.0) or WSL with GPU driver (test set-up: 560.35.05, Nvidia). 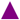 **CRITICAL** For `nicespike`, make sure to have the GPU drivers and CUDA properly installed^38^.
- Software for `nicespike` (0.2.2): CUDA toolkit (test set-up: 12.6), Docker (test set-up: 27.5.1), Git (test set-up: 2.25.1). The docker container that is used includes `python` 3.11.7 with `spikeinterface` (0.100.0)^7^. Further, the modules `nicegui` (test set-up: 1.4.23), `numpy` (test set-up: 1.26.4), `python-neo` (0.13.1)^16^, and `spikeNburst` (0.2.2) are installed during `docker-compose`.
- The source code of `nicespike` is available at Codeberg: https://codeberg.org/spiky/nicespike (doi:10.12751/g-node.yyyyyy).
- Software for `spikeNburst` (0.2.2): `python` (≥ 3.8) with `pip`, may be provided by Anaconda (test set-up: 24.1.2) or from the operating system. Test set-up: With `python` 3.12.6, for reading and exporting the modules `h5py` (3.11.0), `odfpy` (1.4.2), `openpyxl` (3.1.5) are used. The analysis and plotting is done using the `matplotlib` (3.9.2), `numpy` (2.0.1), `pandas` (2.2.2), and `scipy` (1.14.1) modules. The GUI is implemented using `GTK` 3.0 interfaced with `pygobject` (3.48.2). The synchrony measures are calculated using the python module `burst_sync` (2.0.1)^10^ with its dependency on `cython` (3.0.11).
- The source code of `spikeNburst` is available at Codeberg: https://codeberg.org/spiky/spikenburst (doi:10.12751/g-node.xxxxxx).

## Procedure 1: spike sorting and analysis with nicespike

### Install nicespike

- **TIMING < 10 min** The installation of `nicespike` needs to be done only once. An alternative way of installation in a `python-venv` environment is described at https://codeberg.org/spiky/nicespike.
  1. Open a Linux terminal for running the commands in the next step.
  2. Download the code for docker container creation and change into the directory, e.g. by executing

git clone -b docker --single-branch
https://codeberg.org/spiky/nicespike.git
cd nicespike
  3. Select or create an output directory where the analysis results will be stored, e.g. by executing `mkdir output`
  4. Edit the user and group identifier inside the file `docker-compose.yml` to match the used Linux system set-up by modifying the fields `UID` (line 5) and `GID` (line 6) to the ones of the running user. Obtain these identifiers by executing the command `id` inside the terminal.
  5. Edit the volumes inside the file `docker-compose.yml` to include the output directory created in step 3 (first path in line 12) and data input directories (add lines after line 12). 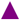 **CRITICAL** Make sure that the user with the identifiers specified in step 4 has read access to the input directories and write access to the output directories specified.
  6. Optionally, edit the host port inside the file `docker-compose.yml` to your preferred (first number in line 10).
  7. Build, compose and run the docker image with:

`docker compose up --detach`
  8. Optionally, close the Linux terminal.

### Open nicespike

- **TIMING < 1 min**
  9. Open the `nicespike` tool in a web browser by accessing http://localhost:8080 (Fig. 2). 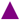 **CRITICAL** If you modified the host port in step 6 you must change it also here. 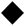 **TROUBLESHOOTING** Alternatively, accessing `nicespike` as a python module is described in BOX 3.

**Fig. 2.**
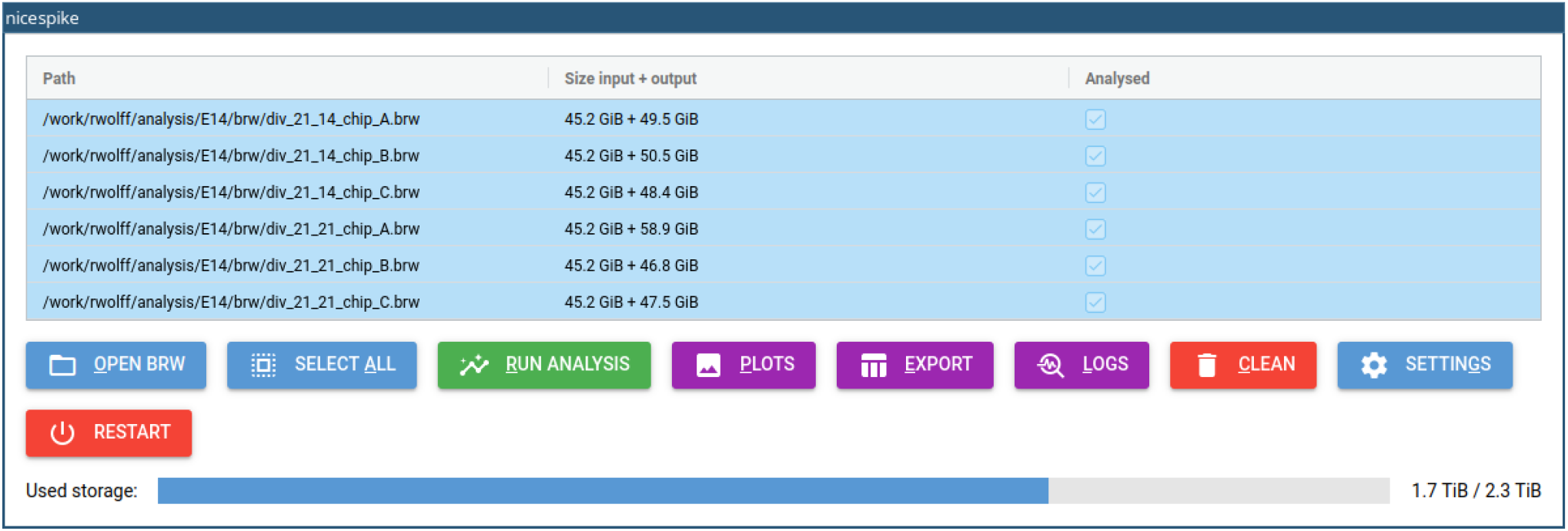
GUI of nicespike after analysing recording files.

### Configure nicespike

- **TIMING < 10 min** The settings may be changed only once or the default settings may be used.
  10. With the opened `nicespike` tool (step 9), open the settings dialog box by clicking the button labelled ‘SETTINGS’. For each setting detailed help is obtained by hovering with the mouse pointer over the setting. See Table 2 for a reference of all parameters that can be set. 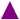 **CRITICAL** Some settings may be hidden by default and scrolling down may be needed to see all.
  11. Adjust the general settings (tab ‘GENERAL’, Fig. 3A, Table 2, General). It is recommended to set a higher number of parallel processes to accelerate the analysis. The export format may be changed, if preferred. The analysis folder should not be changed.
  12. Adjust the settings for spike detection and filtering (tab ‘SPIKES’, Fig. 3B, Table 2, Spikes / Active units). The minimum firing rate is used both for detection and filtering. If the maximum ISI violation ratio is set to a value greater than 0, the units are filtered using the inter-spike-interval violation method^7^. 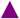 **CRITICAL** Setting the minimum firing rate impacts the template matching. For electrophysiological recordings with very active units, higher minimum firing rates might be appropriate.
  13. Adjust the synchrony settings (tab ‘SYNCHRONY’, Fig. 3C, Table 2, Synchrony).
  14. Adjust the burst detection settings (tab ‘BURSTS’, Fig. 3D, Table 2, Bursts). Depending on the chosen burst detection method, the other parameters below are used or not. The available methods are ‘chiappalone’^19^, ‘pasquale’^20^ (default), and ‘chiappalone_isi_threshold’, which is a modified version of the ‘chiappalone’ method, but using the ISI threshold as calculated within the ‘pasquale’ method.
  15. Adjust the network burst detection settings (tab ‘NETWORK BURSTS’, Fig. 3E, Table 2, Network bursts). The available network burst detection methods are ‘chiappalone’^19^ and ‘pasquale’^20^. Optionally, network bursts can be merged by setting the parameter ‘minimum separation’ to a positive value.
  16. Adjust the plotting settings (tab ‘PLOTTING’, Fig. 3F, Table 2, Plots).
  17. Click on the button labelled ‘OK’.

**Fig. 3.**
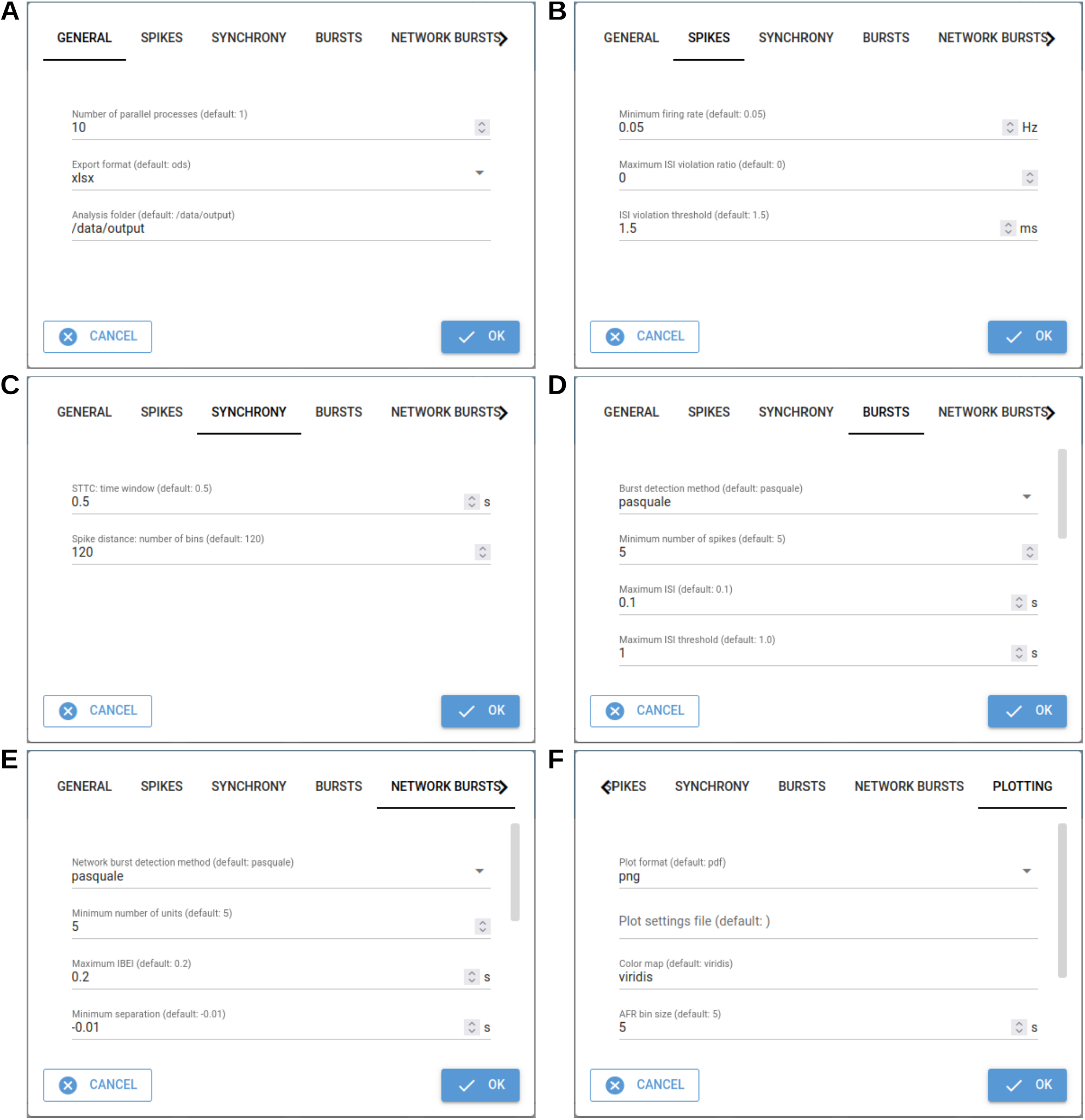
Dialog box tabs for setting parameters in nicespike. (A) General, (B) unit filtering, (C) synchrony, (D) burst and (E) network burst detection, and (F) plotting settings.

### Analyse recordings with nicespike

- **TIMING 30–60 min per recording**
  18. With the opened `nicespike` tool (step 9), open the file loading dialog box by clicking the button labelled ‘OPEN BRW’. Navigate to the folder, where the input files are stored and select either the single `brw` files or folders containing `brw` files. Confirm the selection by clicking on the button labelled ‘OK’. Then, the files are loaded into the application and displayed (Fig. 2). 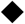 **TROUBLESHOOTING**
  19. Select recordings to be analysed by pressing Shift key on the keyboard and selecting items with the mouse pointer, or select all by clicking on the button labelled ‘SELECT ALL’.
  20. Start the analysis of the selected recordings by clicking on the button labelled ‘RUN ANALYSIS’. This opens a dialog box which allows reviewing the current status of the analysis of the recording that it is currently processed. Cancellation of the analysis will result in restarting `nicespike`. 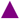 **CRITICAL** In the case if there is no unit found for a recording, an error message is shown, and the concerned recording is ignored in the further analysis. But, there might be other issues, like insufficient storage place (see also the bottom indicator for the used storage, Fig. 2). Thus, it is recommended to study the error messages.
  21. Do plotting of the selected recordings by clicking on the button labelled ‘PLOTS’. Once the plots are collected and archived, a prompt for saving `plots.zip` appears. Choose a folder and save the archive. 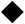 **TROUBLESHOOTING**
  22. Export the selected recordings to tables or python-pickled files (depending on the settings from step 14) by clicking on the button labelled ‘EXPORT’. Once the files are collected and archived, a prompt for saving `export.zip` appears. Choose a folder and save the archive. 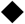 **TROUBLESHOOTING**
  23. Optionally, obtain the log files of the selected recordings for further inspection by clicking on the button labelled ‘LOGS’. Once the log files are collected and archived, a prompt for saving `logs.zip` appears. Choose a folder and save the archive. 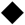 **TROUBLESHOOTING**

### Clean analysed recordings

- **TIMING < 5 min**
  24. To make disk space free, analysis outputs can be cleaned from within `nicespike` by clicking on the button labelled ‘CLEAN’ after selecting recordings (step 19). A dialog box appears with four buttons: ‘CANCEL’ exists the dialog box without doing anything; ‘CLEAN’ erases all analysis output but keeps the files in the table; ‘REMOVE’ drops the selected recordings from the table but keeps the analysis output; ‘DISCARD’ erases the big files from the analysis output but keeps the files (plots, etc.) necessary for exporting. 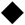 **TROUBLESHOOTING**

Alternatively, analysis outputs, which are stored in the output directory specified in step 3, can be cleaned manually by using operating system tools.

#### BOX 3

**Usage of the `nicespike` as `python` module**

The `nicespike` tool may be accessed via GUI or programmatically within a `python` script or from within a `python` console. The following steps demonstrate its usage with `python` commands.

1. Inside of a python file or a python console, import the `nicespike` tool with

`from nicespike import settings, set_value, SpikeSorting`
2. Optionally, options my be adjusted with the function `set_value`, e.g.

set_value(settings[‘General’][‘Number of parallel processes’], 8)
set_value(settings[‘Spikes’][‘Minimum firing rate’], 0.05)
set_value(settings[‘Network bursts’][‘Minimum number of units’], 5)
set_value(settings[‘Synchrony’][‘STTC: time window’], 0.5)
3. A sorting object is created by

`sorting = SpikeSorting(path=‘brw/div_21_21_chip_A.brw’, output_dir=‘output/div_21_21_chip_A’, settings=settings, verbose=True)`
4. Read the input file and apply the filtering with

`sorting.read_and_filter()`
5. Run the spike sorting using Kilosort with

`sorting.run_kilosort2()`
6. Extract the waveforms with

`sorting.extract_waveforms()`
7. Optionally, export the npz files, which might be used by `spikeNburst` tool, with

`sorting.export_npz()`
8. Run the spike train analysis (burst and network burst detection, synchrony) with
sorting.spikenburst_analysis()
9. Plot and export with

sorting.spikeinterface_plot()
sorting.spikenburst_plot()
sorting.spikenburst_export()

## Procedure 2: spike train analysis with spikeNburst

### Install spikeNburst

- **TIMING < 5 min** The installation of the `spikeNburst` GUI needs to be done only once.
  1. Open a Linux terminal for running the commands in the next step.
  2. A) The recommended way is to create and activate a `conda` environment with

conda create -n spikenburst cython h5py librsvg matplotlib numpy
openpyxl pandas pip python scipy conda-forge::gtk3 conda-forge::odfpy conda-forge::pygobject
conda activate spikenburst B) Alternatively, create and activate a `python-venv` environment with

python -m venv venv_spikenburst
source venv_spikenburst/bin/activate 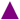 **CRITICAL** It might be necessary to install additional packages for the GUI (GTK 3.0).
  3. Install `spikeNburst` with

pip install --extra-index-url
https://codeberg.org/api/packages/spiky/pypi/simple/ spikenburst[full]

### Analyse spike trains with spikeNburst

- **TIMING 5–20 min per recording**
  4. Open a Linux terminal.
  5. Depending on the choice in step 2 activate the `conda` or `python-venv` environment with
    A. either activate the `conda` environment with

`conda activate spikenburst`
    B. or activate the `python` environment with

`source venv_spikenburst/bin/activate`
  6. Run the `spikeNburst` GUI with

`spikenburst` In all parts of the GUI, hovering with the mouse will show hints for buttons, settings, recordings, etc. 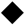 **TROUBLESHOOTING**
  7. Load input data by clicking on the button on the top left with a symbol for opening files (e.g. ‘↥’). A dialog box for file selection appears. Navigate to the folder, where the input files are stored and select either the single `npz` or `bxr` files or folders containing those files. Confirm the selection by clicking on the button labelled ‘OK’. Then, the files are loaded into the application and shown in the table (Fig. 4).
  8. Select recordings to be analysed by pressing `Shift` or `Ctrl` key on the keyboard and selecting items with the mouse pointer.
  9. Select active units by clicking on the button labelled ‘Active’. In the dialog box choose the settings for active units (Table 2, Spikes / Active units, Fig. 5A) and click on the button labelled ‘OK’. The columns labelled ‘Active’ and ‘Spikes’ will be updated with the number of active units and total number of spikes, respectively. 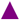 **CRITICAL** Make sure to select proper settings. In particular, the choice, whether spike-sorted units shall be used, depends on the type of input files used.
  10. Optionally, run spike synchrony estimates by clicking on the button labelled ‘Synchrony’. Depending on the chosen method, the parameters can be adjusted (Table 2, Synchrony, Fig. 5B). To use parallelization and speed up the process, increase the number of parallel processes. 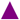 **CRITICAL** If multiple measures of synchrony are to be investigated, repeat the step for each method. 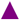 **CRITICAL** For recordings with many (∼ 1000) active units the calculation of synchrony measures can take very long or may fail to complete. Choose STTC for those recordings.
  11. Optionally, detect bursts by clicking on the button labelled ‘Bursts’. In the ‘Burst detection’ dialog box, select proper settings. Depending on the chosen method, the parameters can be adjusted (Table 2, Bursts, Fig. 5C). After clicking on the button labelled ‘OK’, wait some seconds for the analysis completion.
  12. Detect network bursts by clicking on the button labelled ‘Network bursts’. This can be only done if step 11 was run before. Adjust parameters like in step 11 (Table 2, Network bursts, Fig. 5D).
  13. Optionally, plot summary figures by clicking on the button labelled ‘Plot’. Adjust parameters (Table 2, Plotting). 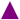 **CRITICAL** Make sure to select a proper location for the output files and the preferred plot format. 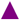 **CRITICAL** The option ‘Plot settings file’ should usually be kept empty. It can be used to specify advanced settings (see https://codeberg.org/spiky/spikenburst/src/branch/main/plot_settings.txt).
  14. Optionally, export the results into tables or python-pickled files by clicking on the button labelled ‘Export’. Adjust parameters (Table 2, General). 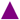 **CRITICAL** Make sure to select a proper location for the output files and the preferred export format.
  15. Optionally, compare different recordings depending on the conditions ‘Cond. A’ and ‘Cond. B’ that can be modified in the columns of the table in the GUI. 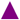 **CRITICAL** This is an experimental feature that produces some comparison plots. It is recommended to use the exported table files from step 14 to produce plots and calculate proper statistics for publication. Alternatively, accessing `spikeNburst` as a python module is described in BOX 4.

**Fig. 4.**
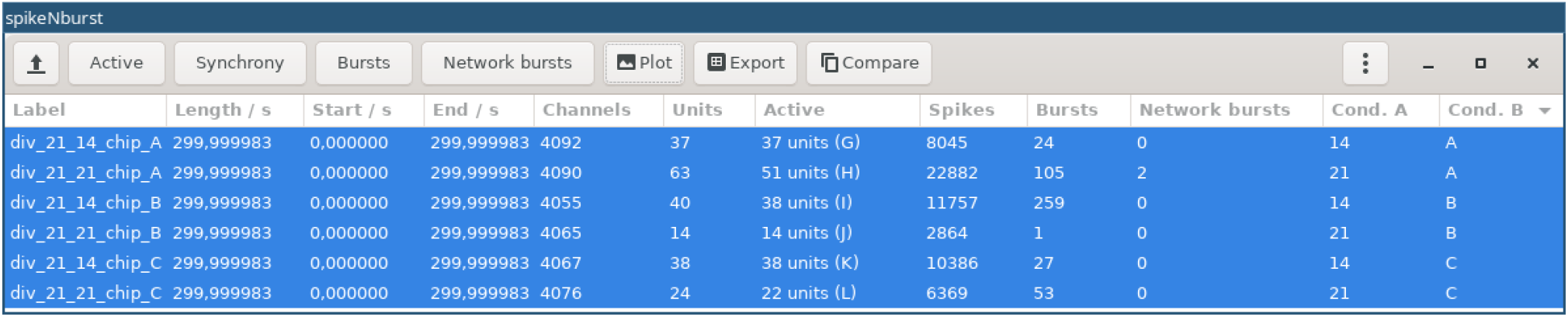
GUI of spikeNburst after analysing recording files.

**Fig. 5.**
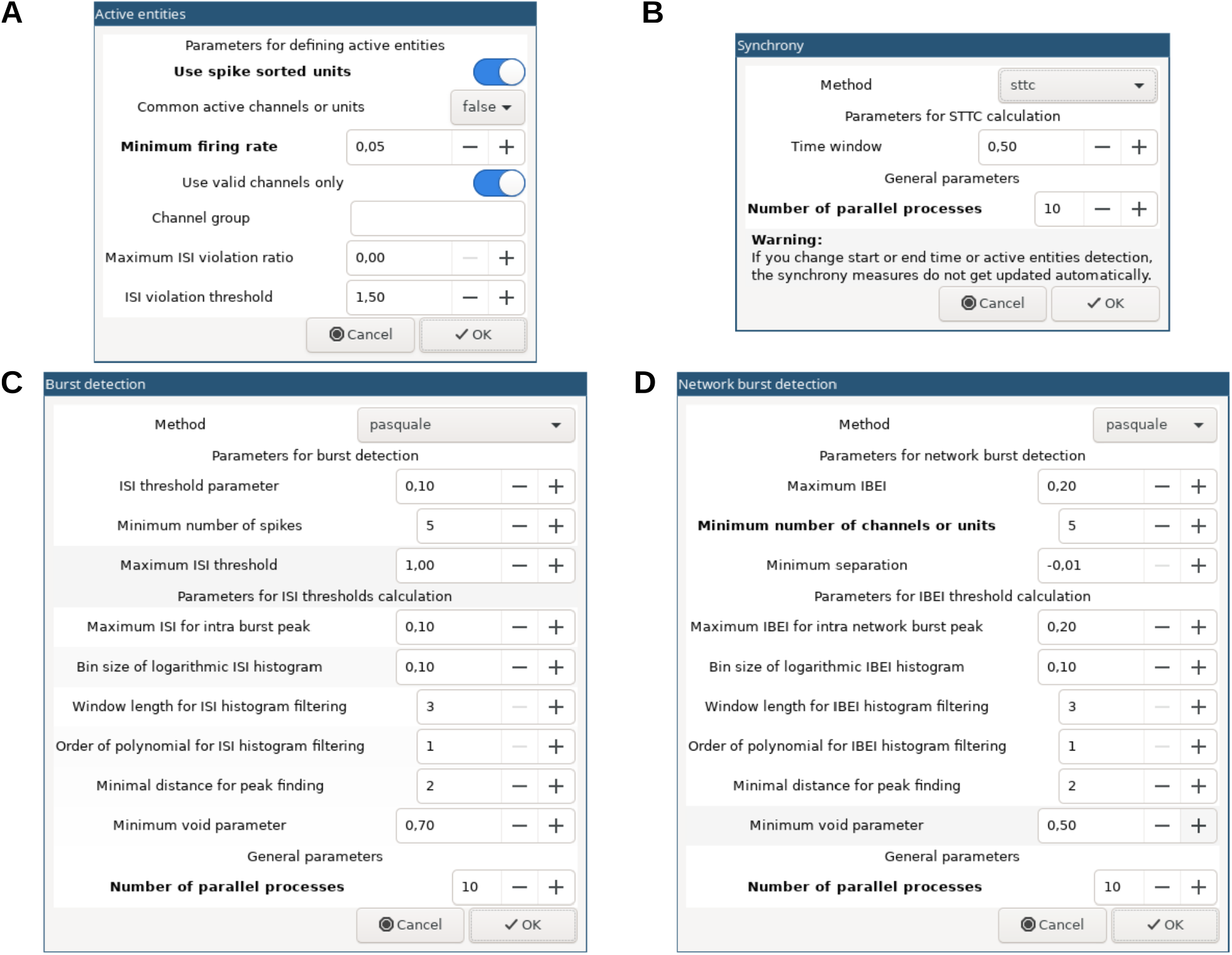
Dialog boxes for setting parameters in spikeNburst. (A) Active units, (B) synchrony, (C) burst and (D) network burst detection settings.

#### BOX 4

**Usage of the `spikeNburst` as `python` module**

The `spikeNburst` tool may be accessed via GUI or programmatically within a `python` script or from within a `python` console. The following steps demonstrate its usage with `python` commands.

1. Inside a python file or a python console, import the `spikeNburst` tool with

`import spikenburst`
2. Create a spike analysis object, specifying parameters by

`sa = spikenburst.SpikeAnalysis(‘npz/div_21_21_chip_A.npz’, use_spike_sorted_units=True, min_firing_rate=0.05)` Available parameters for all classes and functions are documented in the help text, e.g. accessible by

`help(spikenburst.SpikeAnalysis)`
3. Calculate various synchrony measures with

sttc = sa.sttc(dt=0.5)
phase_synchrony = sa.phase_synchrony()
spike_distance = sa.spike_distance(n_bin=120)
4. Run the burst detection on the spike analysis object created in step 2 with

`ba = sa.burst_detection(‘pasquale’)`
5. Run the network burst detection on the burst analysis object created in step 4 with

`nba = ba.network_burst_detection(‘pasquale’, min_n_entity=5)`
6. Plot the results with given prefix

sa.plot(‘ouput/div_21_21_chip_A/s_’)
ba.plot(‘ouput/div_21_21_chip_A/b_’)
nba.plot(‘ouput/div_21_21_chip_A/n_’)
7. Export the analysis results from the network burst analysis object with

`nba.export(‘ouput/div_21_21_chip_A.ods’)` This will also export the results from the spike and burst analysis.

### Troubleshooting

Troubleshooting advice can be found in Table 3.

**Table 3.** Troubleshooting table.

| Step | Problem | Possible reason | Solution |
| --- | --- | --- | --- |
| **Procedure 1** |  |  |  |
| 9 | Connection error | The docker container is not running or the specified port is wrong. | Check that the docker container is running or restart it if necessary, e.g. by executing `docker restart nicespike`<br>Verify that the specified port in step 6 matches the port in the URL. |
| 18 | Input files not found | The folder with the input files is not mounted to the docker container. | Check that the input folder is correctly mounted within the docker container (step 5). |
| 21, 22, 23 | No save-to dialog box | Sometimes, in particular for remote connections, the archive is created, but the user is not prompted to save it. | Check manually (not from within nicespike) the contents of the in step 3 selected output directory and search for a file with name 'plots.zip', 'export.zip' or 'logs.zip'. |
| 21, 22 | Selected recording not in archive | The recording was not analysed or there was an error, or no units were found. | Check if the third column ('Analysed') in the table of the tool for the recording has a tick. If not, run the analysis on that recording (step 20). If there is an error message appearing, check the logs for this recording (either within the error message or using step 23 on this recording). |
| 21 | Missing expected plots | Some type of plots may be disabled. | Check the plotting settings (step 16). |
| 24 | Unresponsive GUI | Deletion of many big files takes time. | Wait some time.<br>Restart the docker container (see troubleshooting of step 9). |
| **Procedure 2** |  |  |  |
| 6 | GUI not shown | Some dependencies are missing, e.g., GTK. | Install the missing dependencies. It is recommended to use an Anaconda environment for their installation. |
|  | Executable not found | The environment was not activated. | Activate the conda or python-venv environment (step 5). |
|  |  | The installation was not successful. | Make sure that the installation inside the conda or python-venv environment completed successfully (steps 1–3). |

### Timing

#### Procedure 1

Steps 1–8, Install `nicespike`: < 10 min

Step 9, Open `nicespike`: < 1 min

Steps 10–17, Configure `nicespike`: < 10 min

Steps 18–23, Analyse recordings with `nicespike`: 30–60 min per recording

Step 24, Clean analysed recordings: < 5 min

#### Procedure 2

Steps 1–3, Install `spikeNburst`: < 5 min

Steps 4–15, Analyse spike trains with `spikeNburst`: 5–20 min per recording

### Anticipated results

Our analysis pipeline, combining `spikeNburst` and `nicespike`, was evaluated on datasets recorded from mESCs-derived neuronal cultures, demonstrating its effectiveness in quantifying synchrony and burst dynamics at both unit and network levels and identifying somatic and dendritic features of single neurons.

### Neurons are recorded from multiple electrodes

We recorded raw voltage traces from wild-type (E14) cells at two (DIV 21+14) and three (DIV 21+21) weeks after the end of the cortical differentiation (BOX 1). We analysed the raw recordings with the described tools using two types of spike detection: 1. the single-channel spike detection from `BrainWave`, and 2. spike-template matching with `Kilosort` within the `nicespike` tool. Subsequently, we performed spike, burst and synchrony measurement using `spikeNburst`. We observed that the mean firing rate was over-estimated by the single-channel spike detection by one order of magnitude (Fig. 6A).

**Fig. 6.**
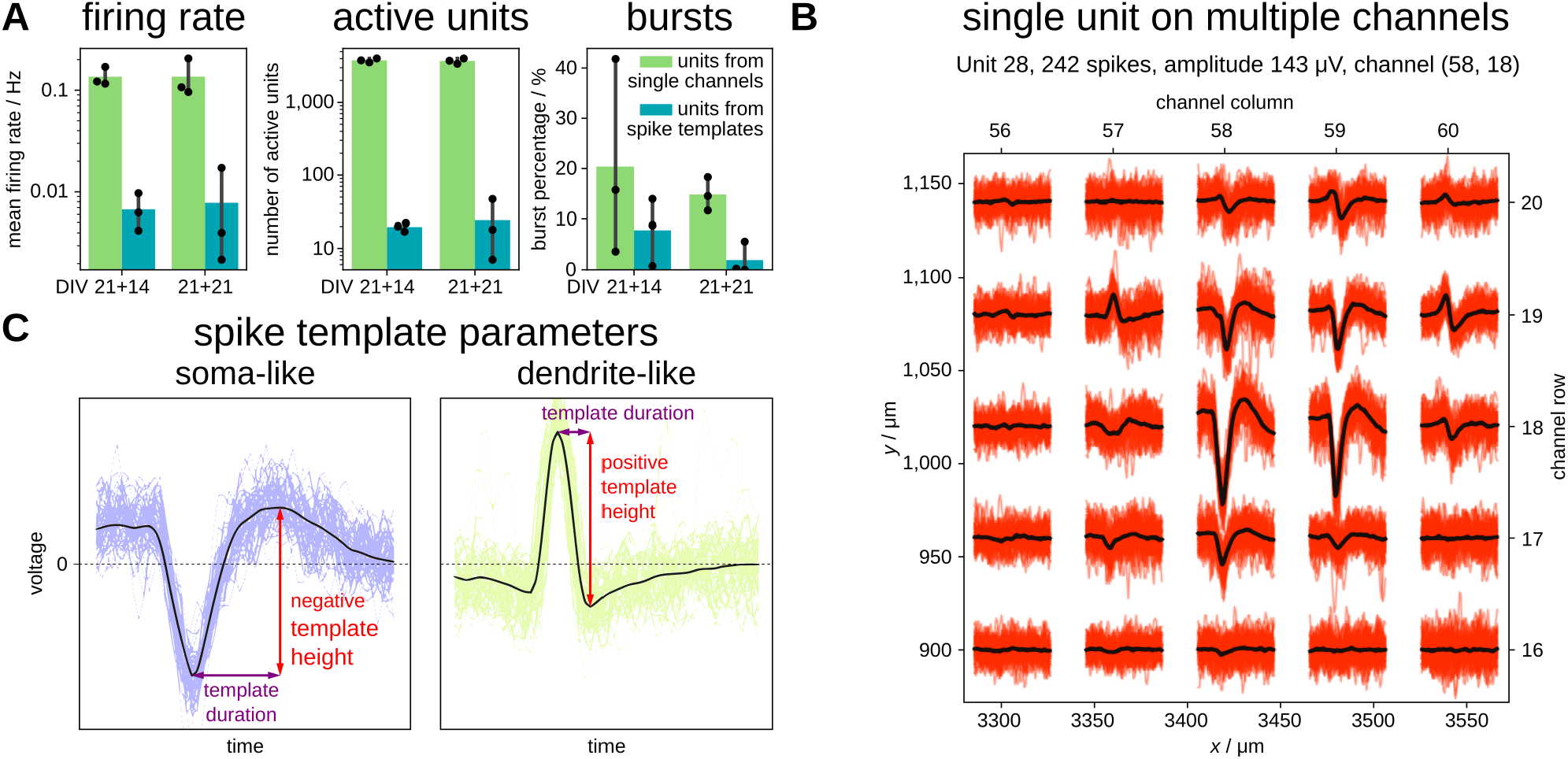
Spikes for single channels and for units from spike-template sorting. (A) Parameters of recordings in three recordings at DIV 21+14 and three recordings at DIV 21+21 using units from single channels and spike template sorting. (B) An example unit that has a matched template with somatic (negative voltage peak) and dendritic (positive voltage peak) parts spanning over at least 200 μm. (C) Estimate of spike template height and duration for soma-like and dendrite-like units.

Duplicates are expected for single-channel spike detection, because the activity of the same neuron may spread on multiple channels. By contrast, the sorting by spike-template matching characterized the neuron’s spatial footprint, identifying the highest spiking activity of a single unit captured by a central channel and the traces of spike activity on the surrounding channels ( Fig. 6B). Also, the number of active units was over-estimated by the single-channel spike detection by two orders of magnitude (Fig. 6A), and more bursts were detected for the units from single-channels than from spike-template matching (Fig. 6A).

We classified units to be soma-like or dendrite-like depending on the spike template shape of the central channel that has the highest voltage difference (Fig. 6B–C). Spike templates where the absolute minimum voltage is greater than the maximum voltage are classified as soma-like with an associated negative template height, while dendrite-like units have a positive template height^27^.

### Automated plotting and exporting with spikeNburst and nicespike

The analysis pipelines implemented both in `spikeNburst` and in `nicespike` include automated plotting and exporting. For each spike-sorted unit the templates are plotted for the central channel and surrounding channels within `nicespike` (Fig. 6B). The plotting with `spikeNburst` includes histograms and spatial plots of many measured parameters (Fig. 7).

**Fig. 7.**
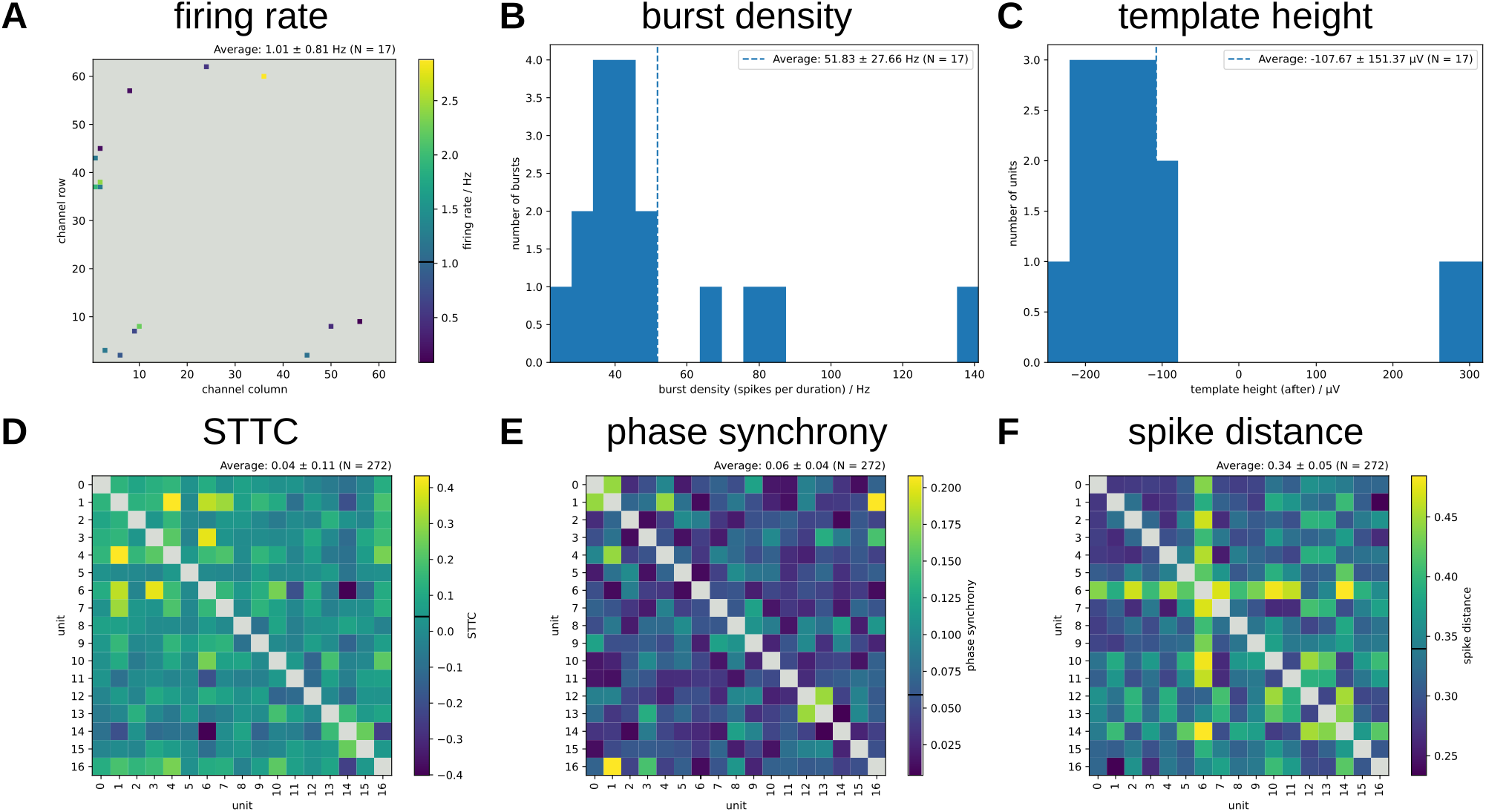
Plots for one recording after spike-template sorting by **spikeNburst**. (A) Firing rate of units at their locations. (B) Burst density of 17 bursts. (C) Unit template height of 2 dendrite-like and 15 soma-like units. (D–F) Various synchrony measures for pairs of units: STTC with Δ*t* = 0.3 s (D), phase synchrony (E) and spike distance with *N*_bin_ = 120 (F). The recording analysed here is for chip A, DIV 21+14 with 17 active units. (A–C) *N* = number of units, (D–F) *N* = 2 × number of unit pairs.

The firing rate of units depending on their location shows that few units have high firing rate of up to around 3 Hz (Fig. 7A). The bursts have a mean burst density of 52 ± 28 spikes per burst (Fig. 7B). Most found units were classified as soma-like, and only two were found to be dendrite-like with a template height of about 300 μV (Fig. 7C). Different synchrony measures have been calculated (Fig. 7D–F). They have different meanings: the STTC and the phase synchrony are 0 for no synchrony and 1 for high synchrony, while the spike distance is 0 for highest synchrony and greater 0 for low synchrony. In particular, the phase synchrony and the spike distance show anti-correlation as expected. Overall, the example recording does not have high synchrony.

In summary, `spikeNburst` and `nicespike` represent a step forward in the field of *in vitro* and *ex vivo* electrophysiology, enabling researchers to efficiently analyse complex HD-MEA datasets and uncover detailed neuronal dynamics. By addressing critical gaps in current analytical approaches, these tools lay the groundwork for future innovations in the study of neural systems and their applications in neuroscience and bioengineering.

## Author contributions

R.W., A.P. and V.T. developed the protocol. A.P. performed the experiments. R.W. implemented the software and algorithms. A.P.B. contributed to the software for the processing of input files from raw electrophysiological recordings. M.C. and V.T. supervised. R.W., A.P. and V.T wrote the manuscript with contributions from A.P.B. and M.C. R.W. and A.P. contributed equally.

## Acknowledgements

Funded by the European Union – NextGenerationEU and by the Italian Ministry of University and Research (MUR), National Recovery and Resilience Plan (NRRP), Mission 4, Component 2, Investment 1.5, project “RAISE – Robotics and AI for Socio-economic Empowerment” (ECS00000035). V.T. and M.C. are part of RAISE Innovation Ecosystem.

